# Genome-wide mapping of gene essentiality in *Pseudomonas chlororaphis* ATCC 9446 using transposon mutagenesis

**DOI:** 10.64898/2026.03.12.711240

**Authors:** Andrés de Sandozequi, Miguel Ángel Bello-González, Omar Alejandro Aguilar-Vera, José Utrilla

## Abstract

Pseudomonas chlororaphis ATCC 9446 is a non-pathogenic rhizobacterium with biotechnological relevance as a biocontrol agent and a promising chassis for synthetic biology. Understanding which genes are strictly required for survival is fundamental to both bacterial physiology and chassis engineering. Here, we generate a genome-wide map of genetic essentiality for P. chlororaphis using high-density Random Barcoded Transposon Mutagenesis (RB-TnSeq). To convert gene-level annotation into biological insight, we layered functional assignments from complementary annotation pipelines, integrating orthology/domain classifiers, ontology mapping, protein export, and delineation of secondary metabolism. These overlays reveal that essentiality concentrates in canonical information processing, envelope biogenesis, and central energy/cofactor and nucleotide metabolism, while large genomic regions are non-essential and therefore represent safe candidates for streamlining and pathway installation. Mapping essentiality onto biosynthetic gene clusters (BGCs) shows that most pathways are dispensable, but some essential genes co-localize, clarifying boundaries for safe editing around BGC loci. Comparison of experimentally determined essential genes with in silico predictions across additional P. chlororaphis genomes show strong overall agreement. Conversely, 32 essential gene orthogroups were found to be conserved across most genomes, yet were classified as non-essential by a prediction algorithm. Together, the resolved essential genome and its integrative functional interpretation provide a durable reference for P. chlororaphis biology and a functional blueprint that can be leveraged for rational streamlining in agricultural, biocontrol and industrial biotechnology.

**Data summary:** The complete genome sequence of Pseudomonas chlororaphis subsp. chlororaphis ATCC 9446 is publicly available in NCBI under RefSeq accession NZ_CP144767.1 (BioProject PRJNA224116; BioSample: SAMN39889236; assembly GCF_036689615.1). Gene annotation, essentiality calls, insertion mapping outputs, and gene-level insertion statistics are provided in Supplementary Table S1. Predicted biosynthetic gene clusters were identified using antiSMASH v8.0 (bacterial version; https://antismash.secondarymetabolites.org); an annotated GenBank file including BGC coordinates is provided as Supplementary File 3 and a summary of BGC features is provided in Supplementary Table S2. Signal peptides were predicted using SignalP 6.0 (https://services.healthtech.dtu.dk/service.php?SignalP-6.0). Protein-coding sequences were functionally annotated using eggNOG-mapper v2 with the eggNOG 5.0 database (https://github.com/eggnogdb/eggnog-mapper), producing Gene Ontology terms and COG functional classifications. Additional functional annotations, including SEED Subsystems classifications, were obtained from BV-BRC (https://www.bv-brc.org/view/Genome/333.24) and are included in Supplementary Table S1. Transposon insertion processing was performed using the FEBA pipeline (PoolStats.R and associated scripts; https://bitbucket.org/berkeleylab/feba), and essential gene inference was performed using the Bio-Tradis pipeline (https://github.com/sanger-pathogens/Bio-Tradis). Comparative essentiality predictions were obtained using DELEAT (https://github.com/jime-sg/deleat). Raw sequencing reads of transposon-genome junctions are available in the NCBI Sequence Read Archive (SRA) under BioProject accession PRJNA1436027.

## Introduction

Defining which genes are indispensable for growth under defined laboratory conditions is a cornerstone of bacterial functional genomics [1–3]. Despite the large and growing number of available genome sequences, a substantial fraction of bacterial genes still lacks definitive functional assignments [4], and for many non-model species we lack precise definitions of the essential genome that link genotype to phenotype. High-throughput pooled transposon mutagenesis coupled to deep sequencing has transformed this field by enabling genome-wide interrogation of loss-of-function phenotypes at scale [4–6]. However, outside a limited set of model organisms, comprehensive descriptions of the essential genome remain sparse, and even curated resources such as DEG [7] only cover a fraction of bacterial diversity. Moreover, the absence of a robust comparative framework across related species hampers our ability to distinguish core, conserved essential functions from clade- or niche-specific requirements [8].

*Pseudomonas chlororaphis* is a non-pathogenic, rhizosphere-associated bacterium noted for its metabolic versatility and production of bioactive secondary metabolites, features that make it attractive for biotechnology and as an emerging chassis for synthetic biology [9]. Several studies have already explored the potential of *Pseudomonas chlororaphis* strains as production chassis for secondary metabolites and as a biocontrol agent [9–23]. Recent studies in *P. chlororaphis* Qlu-1 and H18 have enabled gram-per-liter production of phenazine-1-carboxamide and 1-hydroxyphenazine through medium optimization and genetic engineering, establishing robust platforms for phenazine derivatives [12,19]. Additionally, *P. chlororaphis* ATCC 9446 has been engineered as a non-pathogenic host for rhamnolipid production via *P. aeruginosa rhl* genes, revealing regulatory crosstalk between native and heterologous systems [24]. Complementing these efforts, a genome-scale metabolic model for *P. chlororaphis* ATCC 9446 has provided a system-level view of its metabolic capabilities, including carbon source utilization, denitrification, and phenazine-1-carboxamide production, and has been used to propose strategies to couple growth, redox balance, and secondary metabolite synthesis [21]. Genome-wide essentiality has been defined for several plant-associated and opportunistic Pseudomonads. In the plant-commensal biocontrol strain *Pseudomonas protegens* Pf-5, a dense transposon mutant library combined with transposon-directed insertion site sequencing (TraDIS) was used to identify the essential genome, which is enriched for fundamental cellular functions, whereas genes involved in nutrient biosynthesis, stress responses, and transport are underrepresented and most essential genes are shared between other species of the *Pseudomonas* genus [25]. Similarly, in *Pseudomonas aeruginosa*, transposon insertion sequencing across multiple strains and media, combined with a statistical classifier, defined a core essential genome and showed that essentiality is strongly influenced by both strain background and growth condition [3]. Furthermore, strain resolved functional genomics has shown that gene essentiality is not invariant within a species, because the requirement for a given enzymatic step can be buffered by accessory gene content, paralog redundancy, transport and salvage capacity, or alternative pathway routing that shifts the growth limiting bottleneck to a different reaction [2,8,26,27]. Under this framework, comparative based *in silico* predictors are expected to generate false positives and false negatives for enzymes embedded in networks with bypass options or dosage sensitive paralog partitioning, even when the underlying function remains critical for growth in specific genetic backgrounds or conditions. These observations support interpreting comparative essentiality calls as conditional and motivate empirical essentiality mapping and multi condition fitness assays to resolve lineage dependent constraints [2]. Together, these studies show that essential genomes in *Pseudomonas* are shaped by lifestyle, strain variation and environment, and highlight the need for comparable essential genome definitions in non-pathogenic, rhizosphere-adapted species such as *P. chlororaphis*.

In this study, we defined the essential genome of *Pseudomonas chlororaphis* ATCC 9446 (from here referred to as Pc9446) using a high-density, randomly barcoded transposon insertion mutant library. To interpret these data in a biologically coherent way, we integrated functional annotations from multiple *in silico* pipelines, including orthology- and domain-based annotation, gene ontology, subcellular localization, protein export, and secondary metabolism gene cluster predictions. This integration allowed us to classify individual essential genes into cellular modules and defined functional contexts. In doing so, we obtained a description of the essential genome and its organization into core requirements for growth. In parallel, we delineated genomic regions that are dispensable and therefore represent safe targets for metabolic pathway engineering and genome streamlining.

## Methods

### Strains and Media

The experiments were carried out using *Pseudomonas chlororaphis* subsp. *chlororaphis* ATCC 9446 available at the ATCC culture collection (https://www.atcc.org/products/9446). For mutant library construction, cultures were grown in Luria-Bertani (LB) broth at 30°C with shaking at 250 rpm and LB agar at 30°C with kanamycin.

### Genome sequencing

We generated a hybrid dataset combining short and long reads of *P. chlororaphis* ATCC 9446 genome. Short-reads were taken from Moreno-Avitia 2017. Long-read sequencing was performed on an Oxford Nanopore MinION. Across platforms, aggregate genome coverage was ∼140×. A complete, circular chromosome was assembled *de novo* with Unicycler v0.5.0 in hybrid mode [28], which internally used SPAdes v3.15.5 [29] for short-read assembly and bridged contigs with long reads. Long-read polishing was performed with Racon v1.5.0 [30]. The circular replicon was rotated to place the *dnaA* locus at the origin. The final assembly comprises a single contig representing one chromosome (6,808,083 bp; GC 63%) and no plasmids detected. The assembly is deposited under BioProject PRJNA1074776 and BioSample SAMN39889236. The chromosome is available as NZ_CP144767.1. The corresponding RefSeq annotation ID is GCF_036689615.1 (PGAP v6.10; Apr 5, 2025).

### Generation of the barcoded transposon

The mini-transposon pBAM1 was used as the starting backbone. BAM1 is a Tn5 based element with a reported bias toward GC rich DNA [31]. We modified pBAM1 to incorporate two BsaI sites within the transposon and followed the design of the barcoded Tn5 transposon vector by Wetmore et al., 2015. In this construct, each synthesized 20 nucleotide random barcode is flanked by the constant regions U1 and U2 to enable uniform amplification and sequencing, and the transposon carries a kanamycin resistance cassette. Random barcode oligonucleotides were synthesized by T4 Oligo Corp. (Guanajuato, Mexico) and cloned into the modified pBAM1 by Golden Gate assembly. The resulting barcoded transposon vector library was propagated in electrocompetent *E. coli* λ *pir*+ to obtain high yield supercoiled plasmid DNA and to generate renewable barcode stocks for future use. Prior to introduction into *P. chlororaphis*, the barcode library maintained in *E. coli* was sequenced, confirming approximately 700,000 unique barcodes, with most barcodes represented one to three times in the maintenance pool. Each barcode provides a stable identifier for an individual insertion lineage and enables quantitative tracking of mutant abundance in pooled experiments.

### Generation of the P. chlororaphis ATCC 9446 mutant library

The transposon was delivered into *Pc*9446 WT electrocompetent cells via electroporation. Multiple transformation events were performed using 200 ng of vector per event. Cells were recovered in liquid LB medium for one hour, and aliquots of transformed cells were plated onto solid LB medium supplemented with kanamycin as a selection marker, then incubated for 48 hours at 30°C. After incubation, 100 µL of liquid LB was added to each Petri dish, and all colonies were scraped off with a sterile spatula and transferred to a 3L flask containing LB with kanamycin at a concentration of 50 mg/L, aiming to unify all generated mutants and incubated until the culture reached mid-log phase (0.6 OD_600_). From this point, glycerol stocks of the mutant library were prepared, and samples were saved for genomic DNA extraction and sequencing. Genomic DNA was extracted using the bacterial gDNA isolation kit from Norgen Biotek Corp., following the manufacturer’s instructions for Gram-negative bacteria. Barcode-enriched Tn-Seq libraries were prepared and sequenced by Fasteris Corp. (Switzerland) using their in-house Tn-Seq v6 workflow, which enriches transposon–genome junctions prior to sequencing. Libraries were run on an Illumina MiSeq with single-end 250-bp reads. Mapping of transposon insertions to the genome was done with *MapTnSeq.p*l from the FEBA pipeline [6] and corroborated with Bowtie2 [32]. Reads aligning to more than one genomic location were removed, only uniquely mapped reads were retained for downstream analyses.

### Determination of Essential Genes Using BioTraDIS

To determine the essential genes of *P. chlororaphis* ATCC 9446, we analyzed the transposon insertion data from the mutant library described above using the Bio-Tradis toolkit [33]. Briefly, reads containing the transposon tag that mapped uniquely to the reference genome with high alignment quality (≥98% identity over the first 50 bp adjacent to the transposon end) were retained for further analysis. Gene-wise insertion statistics were obtained with the *tradis_gene_insert_sites.pl* script, which calculates the number and density of insertion sites per annotated coding sequence. Gene essentiality was inferred with the *tradis_essentiality.R* script, which implements an adaptation of the method of Barquist et al., 2016 for TraDIS data. In brief, an insertion index (number of insertion sites normalized by gene length) is computed for each gene, and a loess curve is fit to the genome-wide distribution of insertion indices to identify the minimum between the “essential” and “non-essential” modes. Gamma distributions are then fitted to these two components, and log-odds ratios are used to define an insertion-index threshold for defining genes as essential. Genes with insertion indices below this threshold, including genes with no detected insertions, were classified as essential for growth under our conditions. The resulting essential gene set was subsequently cross-referenced with functional annotations to interpret the biological roles of the essential genome.

### Functional annotation

We annotated all ATCC 9446 proteins with eggNOG-mapper v2 (database eggNOG 5.0) to obtain GO terms (BP/MF/CC) and COG functional categories, with 88% of CDS with assigned COGs [34]. When an annotation for a gene was absent, the information was supplemented with InterProScan domain identification when available [35]. To strengthen and standardize product functions, we cross-validated all functional assignments with the BV-BRC (Bacterial and Viral Bioinformatics Resource Center) RASTtk pipeline, which provided complementary gene functions, EC numbers Subsystem annotations and, when available, additional GO terms [36]. When BV-BRC and eggNOG/InterPro calls disagreed, we prioritized consensus supported by multiple sources, and remaining conflicts were resolved by manual curation using domain composition. To predict and identify N-terminal export signals we used SignalP 6.0 (organism type: Gram-negative; default thresholds) to classify N-terminal signal peptides into Sec/SPI (SP), Sec/SPII lipoprotein (LIPO), Tat/SPI (TAT), Tat/SPII (TAT-LIPO), and pilin-like classes [37].

We tested for over-representation of functional categories among essential versus non-essential genes with one-sided Fisher’s exact tests. *p-values* were adjusted for multiple testing using the Benjamini–Hochberg false discovery rate (FDR), with significance at FDR < 0.05. We report effect sizes as *odds-ratio* with 95% confidence intervals.

To find biosynthetic gene clusters (BGCs), the Pc9446 genome was analyzed with the antiSMASH web server [38] using the Bacteria settings, default parameters, and detection strictness set to “relaxed.” The following auxiliary modules were enabled: KnownClusterBlast, ClusterBlast, SubClusterBlast, MIBiG cluster comparison, ActiveSiteFinder, RRE-Finder (RiPP recognition elements), Cluster Pfam analysis, Pfam-based GO term annotation, TIGRFAM analysis and TFBS analysis. Results were exported as a GenBank file with an updated annotation containing the BGCs information (Supplementary file 3) and a database containing all annotations for each gene is provided as Supplementary Table S1.

### Phylogenomics, Pangenome and Comparative Genomics

We constructed a comparative panel of 85 *Pseudomonas chlororaphis* genomes that included our focal strain Pc9446 and assemblies retrieved from RefSeq that met completeness, quality criteria and were not flagged for contamination. We inferred the species phylogeny with the BV-BRC Bacterial Genome Tree Service [36] using the Codon Tree method on our *P. chlororaphis* genome set (Supplementary Figure S1). The analysis was configured to use 500 single-copy BV-BRC PGFams, enforcing a strict marker set by allowing neither deletions nor duplications. The pipeline estimates a maximum-likelihood tree with RaxML.

To harmonize gene models across the genome set, coding sequences were re-annotated with Prokka v1.14.6, Gram-negative settings [39]. The outputs were parsed into an orthogroup binary presence/absence matrix scored at the genome level, such that an orthogroup was considered present in a genome if at least one gene was assigned to the orthogroup; multi-copy genes were not taken into consideration. Experimentally essential genes were defined exclusively from our RB-TnSeq analysis in Pc9446 and were mapped to orthogroups via the Pc9446 protein identifiers and RefSeq locus tags.

Population structure in the presence/absence matrix was summarized by agglomerative hierarchical clustering of genomes using Jaccard distance with average linkage. The resulting dendrogram was separated into three groups. We calculated prevalence across each group as the size-weighted average. Gene orthogroups were considered core when present in at least 98 % of genomes.

## Results

### Genome-wide transposon mutagenesis and mapping of insertion sites in Pseudomonas chlororaphis ATCC 9446.0

A complete genome sequence for Pc9446 was assembled in this work by combining short reads that were previously deposited as a draft genome [21] with Nanopore long reads to construct a closed single contig genome (RefSeq ID: NZ_CP144767.1) using a hybrid assembly pipeline. We constructed a barcoded transposon mutant library using a modified protocol based on Wetmore et al., 2015. A transposon vector was first constructed to contain a kanamycin resistance marker and a randomized 20-nucleotide barcode flanked by universal priming sites. This pre-barcoded vector was then introduced into wild-type *P. chlororaphis* ATCC 9446 via electroporation to generate the complex library. After antibiotic selection, genomic DNA was extracted and barcode-transposon genome junctions were sequenced (Table 1).

**Table 1.** Genome features of *Pseudomonas chlororaphis* subsp. *chlororaphis* ATCC 9446 and insertion statistics.

| Genome Overview |  | RB-TnSeq Mutant Library Statistics |  |
| --- | --- | --- | --- |
| Genome size | 6,808,083 bp | Total usable reads | 4,352,835 |
| Chromosome topology | Circular | Total insertions detected | 458,697 |
| Total genes | 6,220 | Unique insertion sites | 315,221 |
| Protein-coding genes | 6,059 | Unique insertion sites (central 10–90%) | 220,398 |
| RNA genes | 87 | Insertion density | 1 per 21.63 bp |
| Pseudogenes | 74 | Genes with $\geq 1$ central insertion | 5,755 of 6,059 |
|  |  | Mean insertions per hit gene | 53.3 |
|  |  | Median reads per hit gene | 109 |
|  |  | Mean reads per hit gene | 140.1 |
|  |  | Strand bias (same strand as gene) | 50.90% |
|  |  | Genes with unexpected central insertions | 27 |

The library contained 1 insertion every ∼22 bp, with central insertions in 95% of protein-coding genes and a median of 42 insertions per gene. The distribution of insertions across the genome was visualized using a circular genomic map (Fig. 1), which includes gene orientation, insertion density, and inferred gene essentiality.

**Figure 1.**
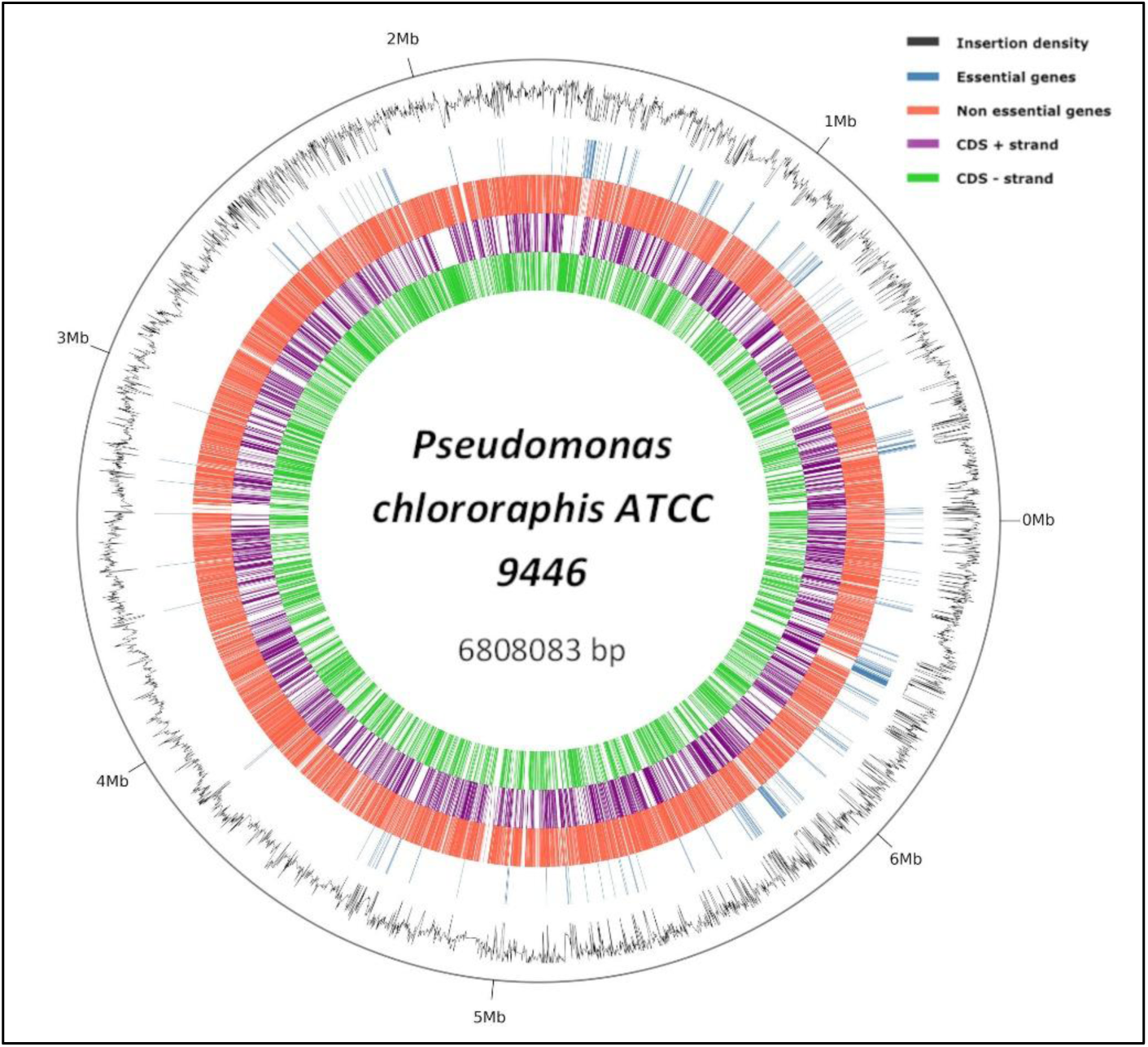
Genome-wide distribution of transposon insertions and essential genes in *Pseudomonas chlororaphis* ATCC 9446. The black line represents insertion density by genomic position, indicating the relative frequency of transposon insertions along the genome. Regions with low or no insertions correspond to genes inferred to be essential in this study. The outermost track shows genome coordinates in megabases.

### Determination of essential genes

To determine essential genes in Pc9446, we employed the Bio-Tradis pipeline [33]. Reads from the barcoded transposon mutant library were mapped to the completed genome of Pc9446. This step generated sorted and indexed BAM files, which were subsequently processed to quantify the number of unique insertion sites per gene and to calculate gene-wise insertion indices. The resulting distribution of insertion indices across genes was distinctly bimodal, indicating the presence of both essential genes, with few or no insertions, and non-essential genes saturated with insertions across the whole length of the gene. For this purpose, we used the Perl script *tradis_gene_insert_sites.pl* to generate gene-level insertion summaries and applied the *tradis_essentiality.R* script, which fits a two-component gamma mixture model to the insertion index data [33]. The essentiality insertion index threshold was set at 0.03209. Genes falling below this threshold were classified as essential (Fig. 2a). A total of 479 genes were classified as essential, taking 7.9 % of all protein coding genes. In 297 essential genes with unique insertion sites (UIS) inside the coding sequence (Fig. 2b), the insertions were not distributed evenly across the coding sequence as in non-essential genes (Fig. 2c) but instead were concentrated near the 3’ end of the gene. This pattern is consistent with insertions being tolerated only when they truncate a small terminal region or fall into segments that are not required for protein activity, whereas insertions within the main coding region disrupt the function and are removed from the population. In contrast, non-essential genes showed insertions broadly distributed across the whole gene length, indicating that disruption of most of the coding sequence is compatible with growth.

**Figure 2.**
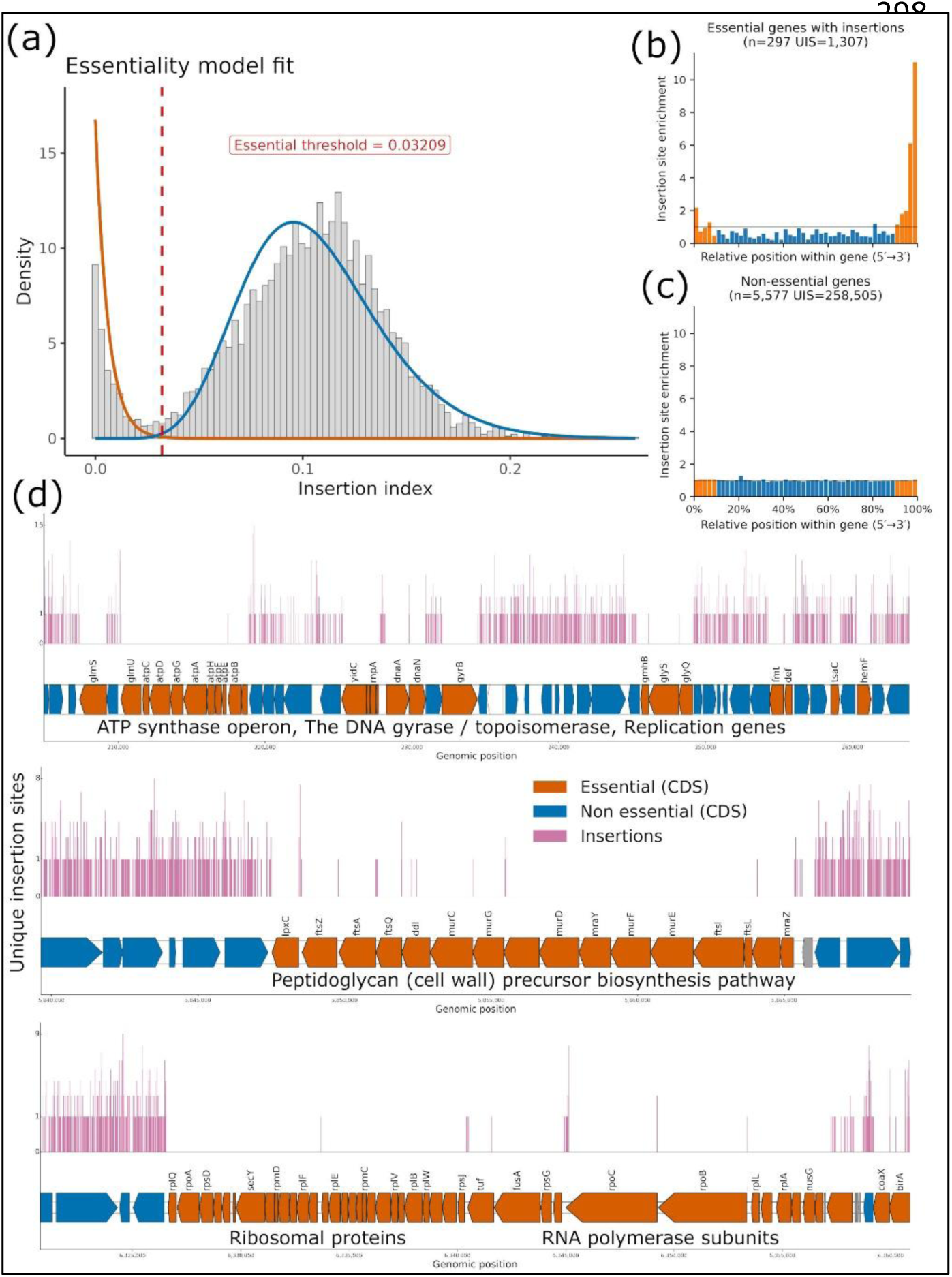
Genomic context of essential and non-essential genomic neighborhoods in *Pseudomonas chlororaphis* ATCC 9446. (a) Genome-wide distribution of normalized insertion indices per gene. The bimodal pattern reflects two gene populations: essential genes with few or no insertions and non-essential genes with high insertion density. Genes with insertion indices below this threshold were called essential. The vertical red line indicates the calculated cutoff. (b) Insertion site profile of essential genes. (c) Insertion site profile of non-essential genes. (d) Representative essential genomic regions. Unique transposon insertion sites are shown along the genome axis to reflect insertion density. UIS, unique insertion sites.

To assess the accuracy of our essentiality assignments, we examined the insertion profiles across genomic regions that are known to harbor essential genes. As expected, we observed a localized depletion of insertions in important loci. These regions exhibited insertion-free zones encompassing core genes such as the ATP synthase operon, replication loci, elongation factors, ribosomal proteins and the RNA polymerase subunits (Fig. 2d). Similarly, the operon involved in peptidoglycan biosynthesis, including *murA*, *murC,* and *mraY*, are located within a genomic island with almost no insertions, highlighting the requirement for cell envelope biogenesis. The sharp contrast between insertion-depleted and insertion-rich regions supports the robustness of the experimental essentiality determination method. Although some genes and intergenic regions between essential loci tolerated insertions, this is indicative of the high degree of insertion saturation of the genome [40–42].

### Functional enrichment, protein subcellular localization, COGs, and Subsystems of essential genes

Functional analysis of the essential gene set (N=479) revealed a strong enrichment for genes involved in core biological processes (Fig. 3). Genes related to translation, including ribosomal proteins and tRNA synthetases, were prominently represented, reflecting the centrality of protein synthesis in bacterial viability. A substantial number of essential genes also contributed to cell wall biogenesis and nucleotide metabolism. Interestingly, a significant fraction of essential genes (65 of 479 essential genes) fell into the “function unknown” category of COG annotations or without any prediction, confirming the fact that many essential bacterial genes remain uncharacterized.

**Figure 3.**
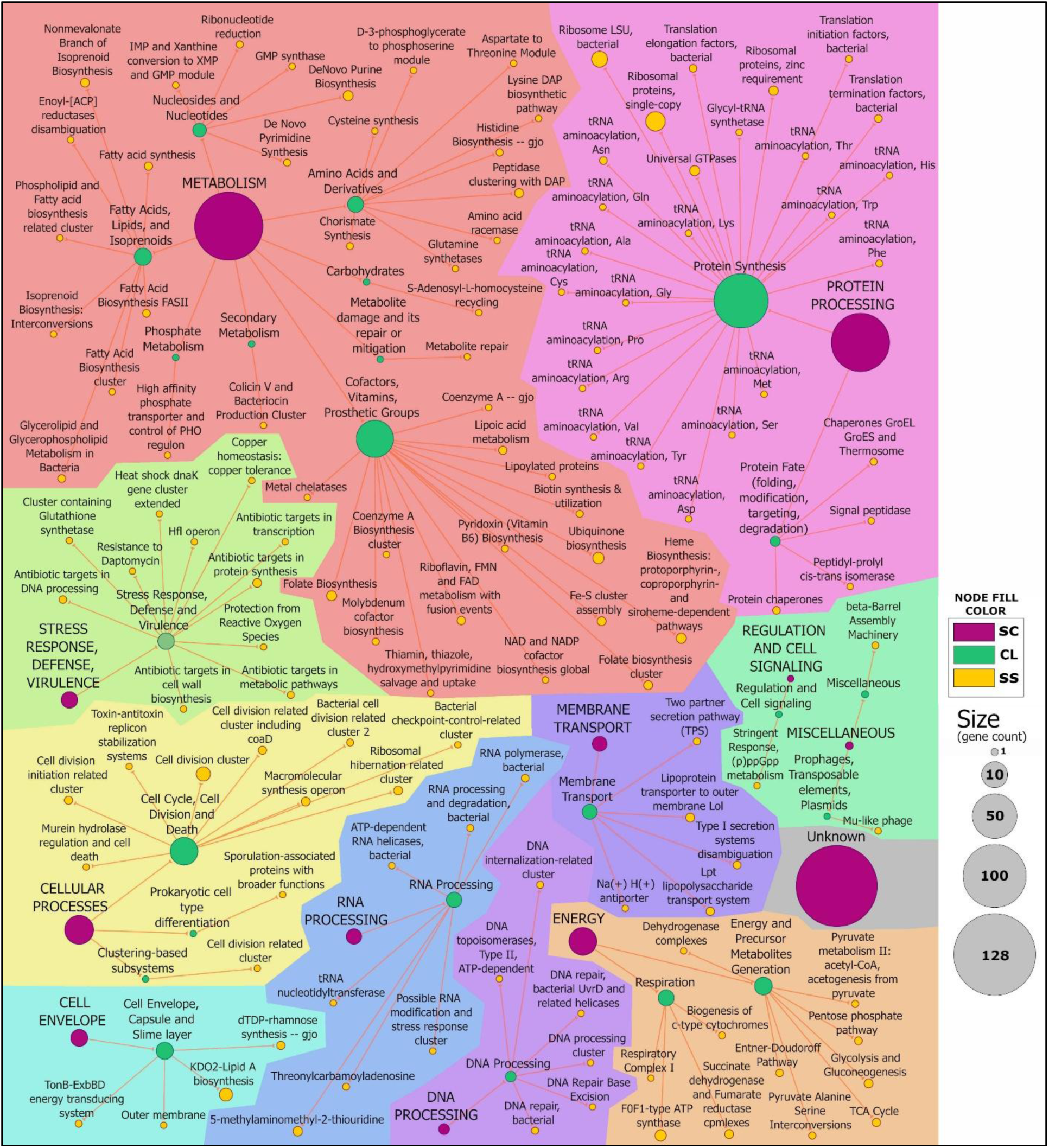
Hierarchical functional network of essential genes. The network represents the SEED-based functional hierarchy linking superclasses (SC) to classes (Cl) and subsystems (SS). Directed edges connect relationships (SC → Cl → SS). Node size is proportional to the number of essential genes assigned to each category.

Beyond these categories, we also identified essential genes in less well-represented pathways. For instance, an ATP-dependent RNA helicase involved in rRNA maturation, an outer membrane protein of a type I secretion system, and enzymes involved in sulfur assimilation and quinone cofactor biosynthesis were all deemed essential. These genes likely contribute to essential functions in ways that are not yet fully understood. Categorical analysis of the essential gene set revealed strong functional enrichments spanning major cellular systems. Based on COG annotations (Fig. 4a), essentials were dominated by translation (COG J; 107 genes, including ribosomal proteins and aminoacyl-tRNA synthetases), followed by a substantial “function unknown” component (COG S; 52 genes), underscoring that many uncharacterized loci are nonetheless required for growth. Additional enrichments were evident for cell wall/membrane/envelope biogenesis (COG M; 49 genes), energy production and conversion (COG C; 43 genes), and nucleotide transport and metabolism (COG F; 41 genes).

**Figure 4.**
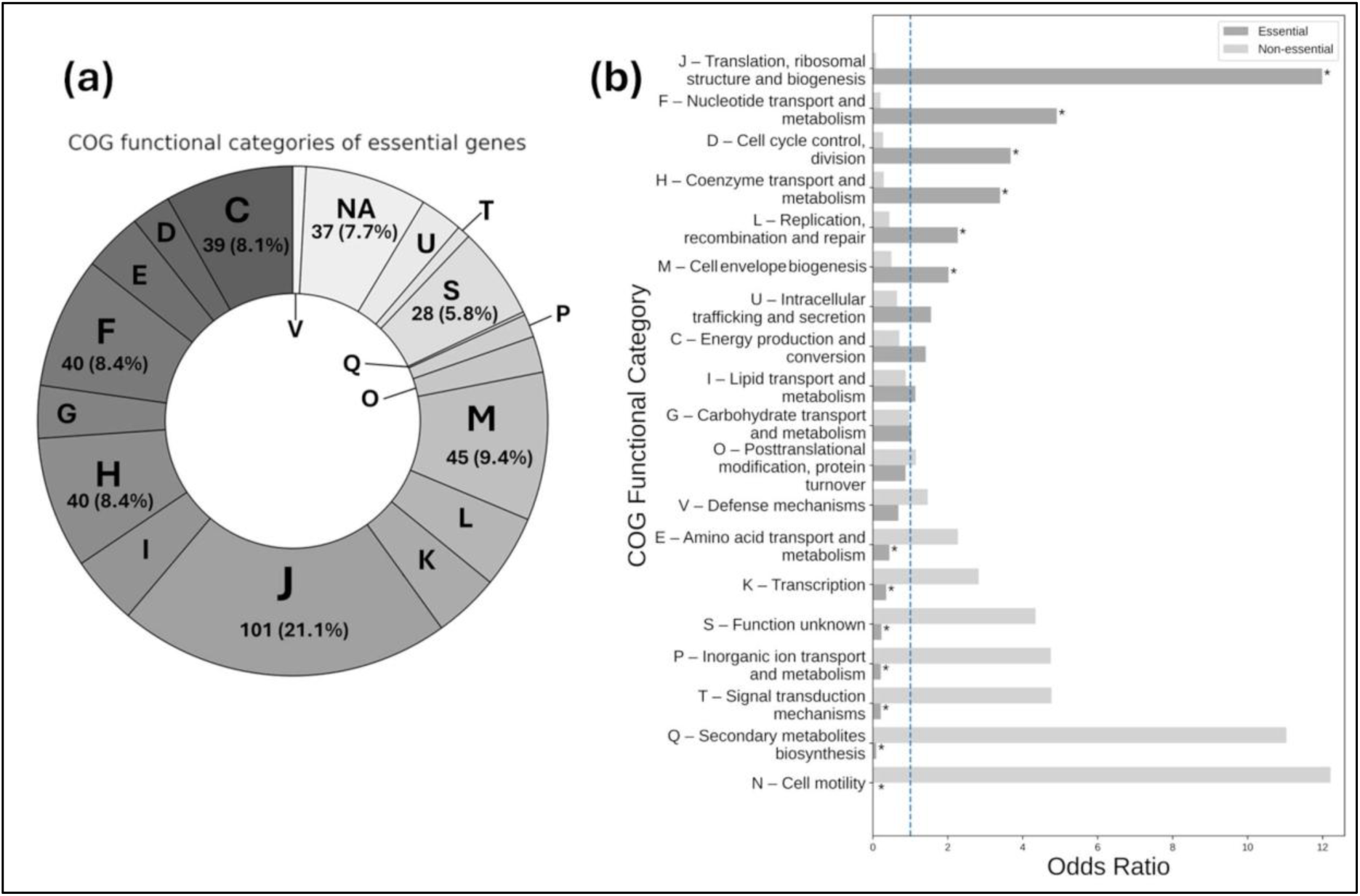
COG functional enrichment of essential and non-essential genes in *Pseudomonas chlororaphis*. (a) proportional distribution of experimentally essential genes across Clusters of Orthologous Groups (COG) functional categories. Each segment represents the fraction of essential genes assigned to a given COG category relative to the total essential gene set. (b) bar plot showing the enrichment of Clusters of Orthologous Groups (COG) functional categories among essential and non-essential genes. Odds ratios were calculated using Fisher’s exact test, comparing the number of genes assigned to each COG category within essential and non-essential gene sets relative to the total genome. A horizontal dashed line at odds ratio = 1.0 indicates no enrichment. Bars represent the odds ratios for each COG category in either essential (dark gray) or non-essential (light gray) gene sets. Asterisks denote statistical significance (*FDR-adjusted P < 0.05*). COG categories A (RNA processing and modification) and B (Chromatin structure and dynamics) were excluded from the plot due to absence of annotated genes in this genome.

At the process level, most essential genes are assigned to ribosome biogenesis (n=45), central metabolism (n=26), RNA processing and modification (n=24), and aminoacyl-tRNA synthetase activity (n=23). The odds ratio analysis was performed to highlight COG categories that were mostly non-essential, that is, enriched in the non-essential gene set (Fig. 4b). These included amino acid transport and metabolism (COG E), transcription (COG K), cell motility (COG N), inorganic ion transport and metabolism (COG P), function unknown (COG S), secondary metabolite biosynthesis, transport and catabolism (COG Q), and signal transduction mechanisms (COG T). Taken together, the essential genome of Pc9446 displays both expected features, such as the inclusion of ribosomal, transcriptional, and cell wall machinery, and a set of intriguing, less-characterized loci that may represent novel essential processes in this bacterium.

### Signal peptide prediction and protein localization

Signal peptides (SPs) are short amino acid motifs that govern protein secretion and translocation across membranes in all domains of life [43]. Although SPs can be predicted computationally, earlier algorithms could not reliably identify the full diversity of signal peptide types. Recent advances, exemplified by SignalP 6.0 [37], apply deep learning to accurately detect all five major classes of signal peptides and can be applied broadly, including to metagenomic datasets. Using SignalP, we identified 851 genes encoding proteins with predicted N-terminal signal peptides, that signal the protein for export to the extracellular milieu via the Sec and Tat pathways or have a lipoprotein signal peptide (Table 2).

**Table 2.** Signal peptide classification predicted by SignalP 6.0.

| Category | Number of proteins (%) | Essential proteins (%) |
| --- | --- | --- |
| No signal peptide (NoSP) | 5,208 (86.7) | 461 (96.9) |
| Sec/SPI (SP) | 582 (9.7) | 13 (2.7) |
| Sec/SPII lipoprotein (LIPO) | 217 (3.6) | 5 (1.1) |
| Tat/SPI (TAT) | 32 (0.5) | 0 (0.0) |
| Tat/SPII lipoprotein (TATLIPO) | 5 (0.1) | 0 (0.0) |
| Pilin-like signal | 15 (0.2) | 0 (0.0) |

Among the functionally annotated cell envelope-related essentials, several correspond to established determinants of outer membrane biogenesis and stability: the outer-membrane lipoprotein LolB, which is required for insertion of lipoproteins into the outer membrane [44]. LolB has been identified as an essential gene for *E. coli* and required for virulence on several phytopathogenic and plant-associated bacteria [45–47]; the LPS-assembly proteins LptD and LptA, central to translocating lipopolysaccharide to the cell surface [48]; the periplasmic chaperone SurA, which assists in folding and maturation of outer membrane proteins (OMPs) [49]; the TolQ component of the Tol–Pal system, involved in maintaining outer membrane integrity [50,51]; the N-acetylmuramoyl-L-alanine amidase AmiC, which participates in cell wall remodeling during cell division [52]; the β-barrel assembly machinery protein BamA, essential for OMP insertion [53]. Uncharacterized essential exported proteins are likely to play critical roles in cell envelope biogenesis, structural maintenance, stress response and gene regulation. Their essentiality, combined with these signal peptide predictions, makes them high-priority targets for functional characterization to uncover novel components of the *Pseudomonas* envelope and physiology.

### Biosynthetic gene clusters

*Pseudomonas chlororaphis* strains are prolific producers of secondary metabolites, including phenazine derivatives and siderophores such as pyoverdine, which contribute to pathogen suppression, iron acquisition, and rhizosphere competitiveness [22,23,54–59]. These metabolic traits underpin their use as a biocontrol agent and make it a valuable model for studying and engineering secondary metabolism [11,13]. To investigate gene essentiality on secondary metabolism loci, we first used AntiSMASH to delineate the boundaries of all BGC in the genome. AntiSMASH predicted 18 biosynthetic gene clusters, similar to a recent study where they predicted 17 BGCs. combined the essential gene set onto the predicted biosynthetic gene clusters (Table S2).

All clusters encompassed 438 unique genes. Of these 18 BGC, 6 of the predicted BGC contained genes that were classified as essential. Essentiality within BGCs was heterogeneous across clusters. Some regions were completely dispensable, whereas others carried a small but non-negligible number of essential genes. For example, the arylpolyene biosynthetic cluster (BGC2), where 2 of 41 genes were essential. The butyrolactone cluster (BGC12) harbored 1 essential among 7 genes. In non-ribosomal peptide clusters, the NRPS region (BGC8) contained 2 essentials in 32 genes. Interestingly, the large pyoverdine BGC (predicted as two separate regions 14 and 15, comprising 72 and 41 genes, respectively) and the phenazine BGC (region BGC17 comprising 22 genes) contained no essentials, consistent with the expectation that iron-acquisition pathways and environment related metabolites are dispensable in laboratory conditions [14]. Essential genes in most BGCs are likely flanking housekeeping genes or envelope biogenesis factors embedded within BGC boundaries. In contrast, the region BGC18 was found to be essential, having 7 essential genes of 24 (Fig. 5).

**Figure 5.**
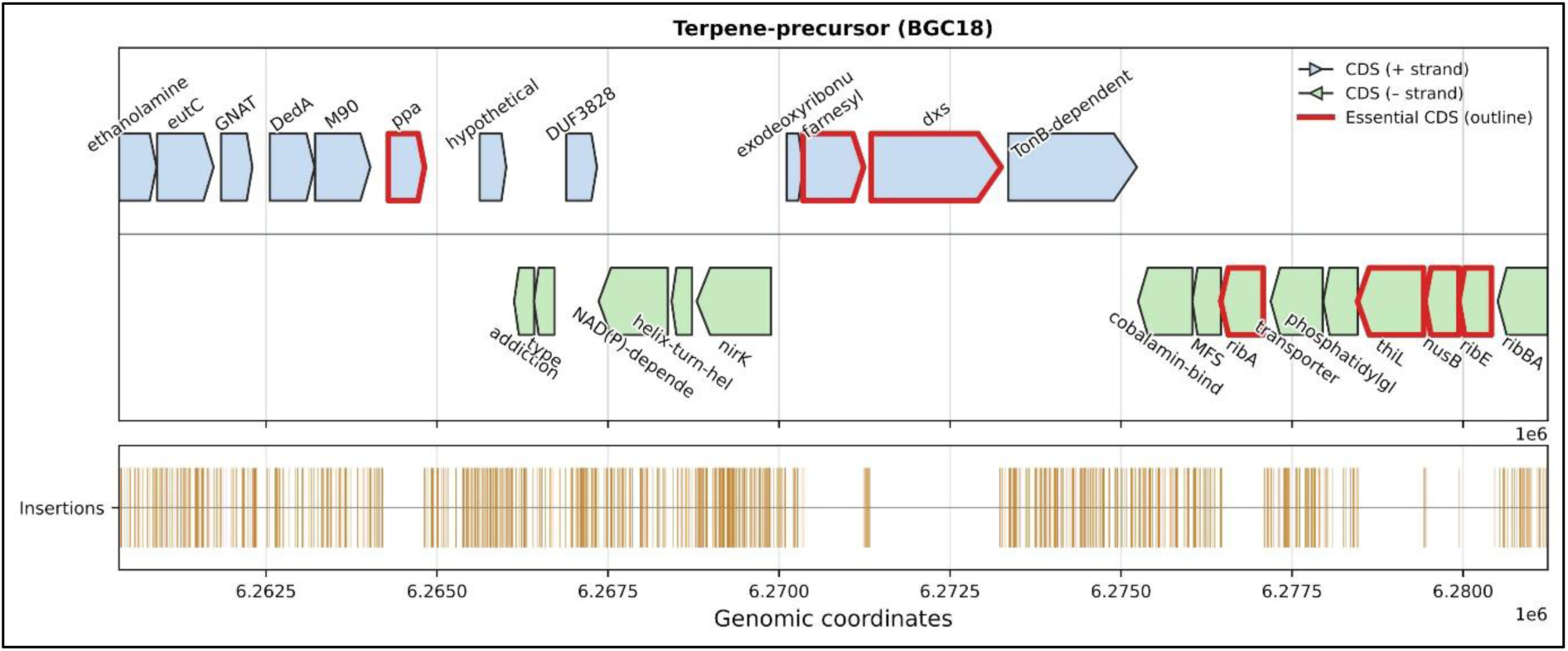
Terpene-precursor (BGC18) map with transposon insertions. Gene organization of BGC18 on the main chromosome (6,260,350–6,281,238 bp). Essential genes determined experimentally by transposon insertion analysis are highlighted with a red outline. All unique transposon insertion sites detected within the BGC interval are shown.

Although BGC18 was annotated as a terpene-precursor cluster, detailed inspection revealed that essentiality (7 essential genes among 24) derives not from sesquiterpene synthesis per se, but from embedded vitamin and isoprenoid precursor pathways. *ribA* and *ribE* anchor riboflavin biosynthesis, *thiL* activates thiamine monophosphate to ThDP, and *dxs* and *ispA* connect to the MEP pathway and FPP production, a precursor also required for ubiquinone side-chain synthesis.

Flanking essential housekeeping genes (*ppa*, *nusB*) further explain the observed signal. Thus, BGC18 functions as a vitamin and cofactor supply hub integrated with isoprenoid precursor metabolism. Across clusters, essential genes within BGC intervals were enriched for housekeeping or envelope-related roles rather than canonical secondary metabolism catalytic domains. Specifically, this cluster anchors two vitamin routes and the isoprenoid precursor pathway: *ribA* and *ribE* correspond to the riboflavin (B2) pathway toward FMN/FAD, while *thiL* carries out the thiamine activation to ThDP (B1), explaining their essentiality. Additionally, *dxs* feeds the MEP pathway from DOXP and, together with *ispA*, establishes flux to FPP, the universal sesquiterpene precursor and a key substrate for ubiquinone side-chain synthesis, which has been characterized as essential in bacteria [60–62]. Flanking this essential, the region carries mostly non-essential functions that likely support cofactor scavenging, redox balance, and local control (a TonB-dependent receptor, an MFS transporter, a cobalamin-binding protein adjacent to *eutC*, *nirK*, a GNAT acetyltransferase, and a helix-turn-helix regulator), as well as maintenance elements such as a toxin–antitoxin pair. Two housekeeping genes, *ppa* (inorganic pyrophosphatase) and *nusB* (transcription antitermination), are also essential, consistent with their biological roles rather than pathway specificity [63]. *ppa* is essential for cell growth in all known organisms and *nusB* has been characterized to be essential in several organisms [64]. Altogether, BGC18 functions as a vitamin/cofactor supply hub intertwined with isoprenoid-precursor generation: it consolidates the *ribA*–*ribE* system for riboflavin, *thiL* for thiamine, and the *dxs*–*ispA* conduit to FPP, thereby linking cofactor homeostasis to sesquiterpene precursor availability [65], explaining the essentiality embedded within an otherwise largely non-essential neighborhood.

### Comparison of experimentally determined essential genes with in silico predictions

To extrapolate our experimental results, we compared our experimentally characterized essential genome to *in silico* predictions from DELEAT [66], a machine learning approach that integrates genomic and functional features to infer essentiality in non-model organisms. Our RB-Tnseq essentiality dataset was compared against DELEAT predictions across 85 *Pseudomonas chlororaphis* genomes organized into orthogroups (Fig. 6a), uncovering a conserved core of 326 essential orthogroups. Experimental–computational agreement was strong overall, with 282 essential orthogroups and predicted by DELEAT to be essential in all other genomes (Fig. 6b, 6c, Tables S3-S5).

**Figure 6.**
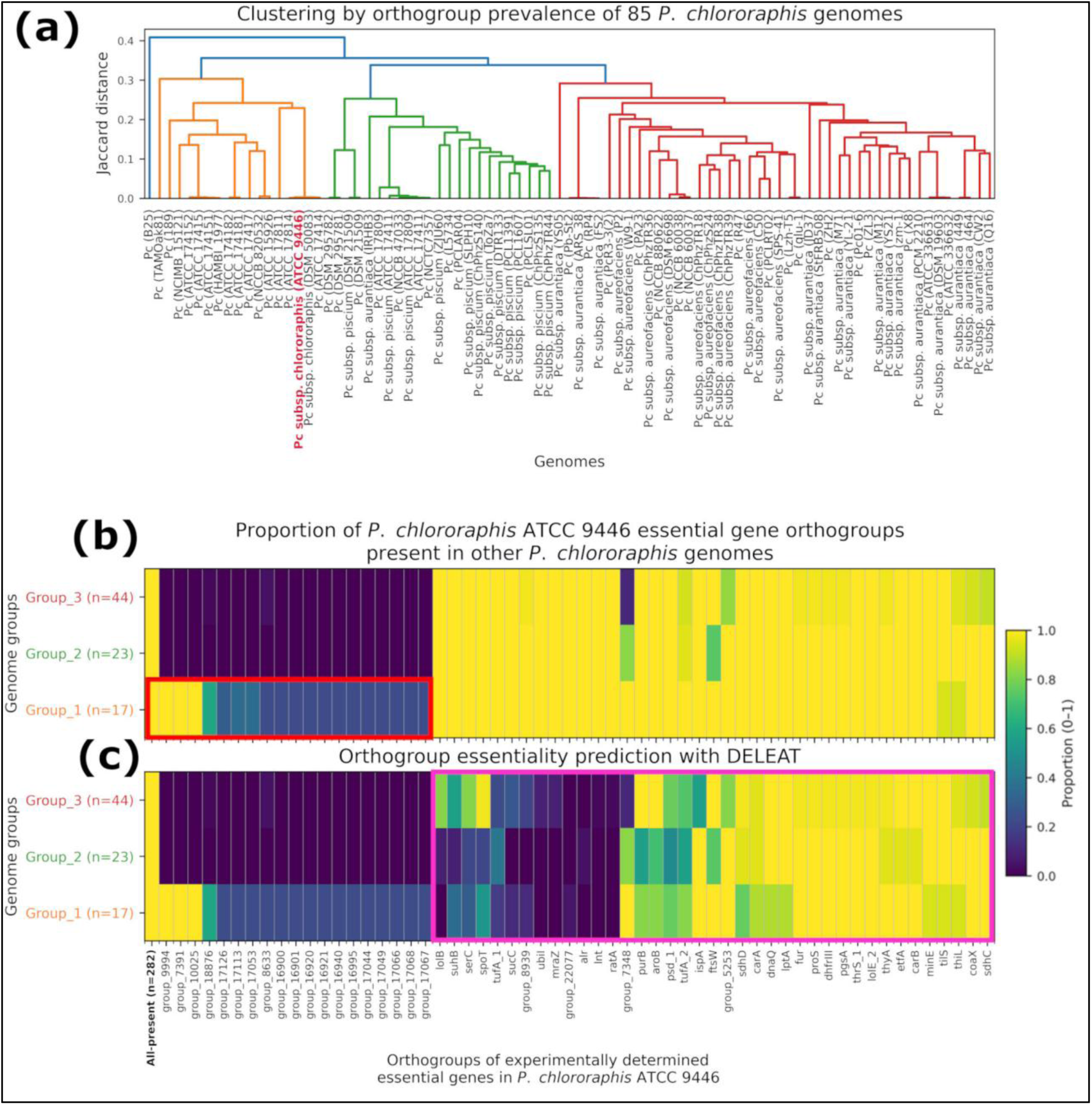
Comparative clustering and group-level essentiality across *P. chlororaphis* genomes. (a) Genome dendrogram. Hierarchical clustering of genomes based on Jaccard distances calculated from the orthogroup presence/absence matrix using average-linkage agglomeration. Leaf labels correspond to strain names as deposited in NCBI. The focal genome analyzed in this study (belonging to group 2) is highlighted in red. Jaccard-based groups are congruent with the inferred phylogeny of the clade (Supplementary Fig. 1). (b) Experimentally essential orthogroups heatmap, for each genome group, the proportion of genomes that contain at least one ortholog to an experimentally determined essential gene in *Pc*9446. (c), benchmarking *in silico* prediction of essential genes in 85 *P. chlororaphis* strains. Using the same column order and set of orthogroups as in panel (b), the heatmap shows, for each genome group, the fraction of genomes in which DELEAT predicts to have an ortholog gene to an essential gene in *P. chlororaphis* ATCC 9446.

A set of 20 essential orthogroups that were found only within the *Pc*9446 closest group of strains, comprised of our *Pc*9446 strain and the strains ATCC 17814, DSM 50083, and ATCC 17414 (Fig. 6b, red rectangle). Parallelly, there were 32 orthogroups of Pc9446 essential genes that were present in most genomes and DELEAT predicted that they were not essential (Fig. 6c, pink rectangle). These include folate biosynthesis enzymes, a thymidylate synthase [67,68], a type-3 dihydrofolate reductase, a type III pantothenate kinase, a thiamine-monophosphate kinas, both copies of the elongation factor Tu-A, LptD for LPS assembly, the peptidoglycan glycosyltransferase FtsW, the Type I secretion membrane-fusion protein PrsE, an RTX-translocation ATPase and the lipoprotein LolB that mediates the outer membrane anchoring of lipoproteins [69–71].

## Discussion

This study establishes a genome-scale, experimentally validated map of genetic essentiality in *Pc*9446 based on a barcoded transposon library. Randomly barcoded transposon sequencing (RB-TnSeq) quantifies gene essentiality by tracking the fate of uniquely tagged insertion mutants during competitive growth [1,6,25,72–80]. These methods provide sensitive, genome-wide characterization of loci required for viability in defined conditions. In parallel, computational approaches can now predict essentiality directly from genome content and comparative signals across many species [7,66,81,82]. When combined, experimental and comparative predictions are complementary: experimental data provides a reliable essentiality assessment, whereas cross-species comparisons reveal whether an essential gene is conserved, duplicated, buffered by paralogs, or restricted to certain lineages or ecological niches [23,81,83].

Using a two-component gamma mixture model applied to insertion indices, we identified 479 essential genes, representing 7.9% of protein-coding loci. The library achieved high genome-wide coverage, and the insertion density firmly sits within the regime of highly saturated libraries suitable for gene-level essentiality inference. Importantly, insertion-free gaps were analyzed considering prior hyper-saturated libraries studies, which demonstrated that short gaps could arise by chance even at extreme densities. The presence of small insertion-free segments in Pc9446 is therefore not interpreted as evidence of essentiality without statistical support. Conversely, complete and extended depletion across loci such as *rpoA*, *rpoB*, *rpoC*, ribosomal proteins, *murA*, *murC*, and *mraY* provides strong biological validation of experimental essential inference. The fact that very short non-essential genes retain multiple insertions, while small but functionally constrained genes show depletion, further indicates that classification reflects biological necessity rather than technical under-sampling.

Functional enrichment mainly serves as a quality and biological coherence check, because the essential gene set concentrates on the expected viability modules, especially translation, envelope biogenesis, nucleotide metabolism, and energy generation (Table S6). This modular structure is consistent with the architecture of essential genomes reported in diverse bacteria, where viability is maintained by a limited number of tightly coupled core systems rather than dispersed single genes. The more interesting observation is the persistent contribution of poorly annotated loci, since a sizeable subset of essential genes remains classified as COG S or unknown even when using subsystem level annotation. In contrast, COG categories enriched among non-essential genes (Fig. 4b) included amino acid transport and metabolism (COG E), transcriptional regulators (COG K), cell motility (COG N), inorganic ion transport (COG P), signal transduction (COG T), and secondary metabolite biosynthesis (COG Q). This pattern aligns with the ecological lifestyle of *P. chlororaphis*, where environmental sensing, motility, and specialized metabolite production contribute to rhizosphere fitness [23] but are dispensable for growth in rich laboratory media.

Signal peptide prediction identified 851 genes encoding exported or membrane-targeted proteins. Among essential genes, multiple envelope-associated factors were required, including *lolB*, *lptD*, *lptA*, *surA*, *tolQ*, *bamA*, and *amiC*, reinforcing the central importance of outer membrane integrity and cell wall remodeling. The presence of essential yet poorly characterized exported proteins highlights the envelope as a key frontier for discovering novel core functions in *Pseudomonas* biology. In addition to well-annotated housekeeping genes, several essential loci encode proteins containing domains of unknown function (DUFs). Some of these correspond to domains previously described as essential DUFs (eDUFs), which are linked to core processes such as ribosome maturation, initiation of DNA replication, and cofactor-related functions [84]. Other DUF-containing essential proteins in Pc9446 (Table 3), were not previously classified as eDUFs. Their essentiality indicates that conserved but still poorly characterized proteins contribute directly to core cellular functions in this organism.

**Table 3.** DUF-containing essential proteins in *Pseudomonas chlororaphis* ATCC 9446.

| Domain<br>Architecture | Annotation | Described as eDUF |
| --- | --- | --- |
| DUF1289 | Predicted Fe–S protein | Yes |
| DUF150 | RimP ribosome maturation factor | Yes |
| DUF177 | YceD family protein | Yes |
| DUF2331 | EarP arginine rhamnosyltransferase | No |
| DUF3203 | DUF3203 family protein | No |
| DUF3470 | Fer4-containing ferredoxin | No |
| DUF493 | Lipoate regulatory protein YbeD | Yes |
| DUF721 | DciA replication initiation factor | Yes |
| DUF5625 | Predicted lipoprotein | No |

These genes provide defined targets for future genetic and biochemical studies aimed at clarifying their roles in *Pseudomonas* physiology and improving functional annotation across the genus. In addition, prophage regions were also mostly non-essential, yet 4 cI-like repressors (locus: V4Y05_RS22805, V4Y05_RS08275, V4Y05_RS11705, V4Y05_RS07010, V4Y05_RS07010) behaved as essential, consistent with the requirement to maintain prophage silencing to prevent lethal induction [85]. In one prophage region, integration near essential genes such as *purC* or *fdxA* further demonstrates how mobile elements can overlap essential loci [86,87].

The essentiality map for Pc9446 is broadly consistent with essential gene sets reported for other plant associated pseudomonads, supporting the existence of a conserved core required for growth in rich medium. In *Pseudomonas protegens* Pf 5, TraDIS identified 446 essential genes and showed the expected enrichment for core cellular processes, with most essential genes belonging to the species core genome [25]. Pc9446 shows a comparable scale and a strong overlap with a conserved orthogroup core, but the comparison with DELEAT highlights why purely comparative inference remains incomplete [2,66,82]. Although overall concordance was high, the discordant orthogroups concentrate in categories that are known to challenge conservation-based prediction, including background dependent essentiality in a pangenome context and cases shaped by redundancy and paralogy [2].

In Pc9446, the 32 orthogroups that were experimentally essential but not predicted as essential in most *P. chlororaphis* genomes are consistent with buffering by alternative salvage or transport capacity, paralog partitioning, or pathway routing that changes which reaction becomes growth limiting. This includes folate and thymidylate associated steps and other cofactor related kinases, as well as envelope and division functions such as *lptD*, *ftsW*, and *lolB*, where requirement can be obscured by annotation differences, paralog substitution, or variation in envelope architecture. The presence of both elongation factor Tu copies in the mismatch set further illustrates a limitation of *in silico* predictions, because essentiality can depend on dosage and regulatory partitioning between paralogs rather than presence alone [2,81,83,88]. The phylogenetic structure of the species also helps explain these discrepancies, as the genomes fall into distinct groups corresponding to *P. chlororaphis* subsp. *chlororaphis*, *P. chlororaphis* subsp. *piscium*, and *P. chlororaphis* subsp. *aurantiaca/aureofaciens* cluster (Supplementary Figure S1). Differences among these groups in accessory functions, pathway context, and cell envelope related traits are likely to reduce the extrapolation of essentiality predictions across the species. Conversely, the 20 orthogroups that were essential only within the closest Pc9446 clade support the idea that lineage restricted gene content and regulatory coupling can create strain group specific dependencies. These patterns should be interpreted alongside the extensive accessory variation in *P. chlororaphis*, particularly across biosynthetic gene clusters and other niche adaptation loci [23].

### Limitations and future directions

Essentiality was determined under a rich-medium condition, and genes classified as non-essential here may become critical under minimal media, carbon and nitrogen sources, rhizosphere-like environments, or bioreactor-relevant stresses [6,12,41,89–92]. Transposon-based approaches also have known limitations, including potential polar effects, sequencing artifacts, redundancy masking, and challenges with very short genes [6,41]. Furthermore, essentiality defines viability thresholds but does not directly quantify metabolic cost or flux redistribution [90].

Genome reduction has been used as a chassis development strategy in *Pseudomonas* to simplify genetic backgrounds and improve secondary metabolite output [14,65,93–96]. In *Pseudomonas chlororaphis* GP72 [15], comparative genomics guided sequential deletions of regions enriched in strain specific DNA and secondary metabolism loci, and some reduced derivatives showed increased phenazine production, although cumulative deletions were also accompanied by changes in colony morphology and a slower growth rate. Our RB-TnSeq essentiality map and the associated barcoded mutant library provide a complementary framework for this type of engineering. Candidate deletions can be selected using empirical dispensability, and barcode-based fitness assays across diverse media and stress conditions can quantify tradeoffs and identify conditionally important genes prior to genome editing. This combination should support genome streamlining in *Pseudomonas chlororaphis* that removes a larger fraction of dispensable sequence with more predictable effects on growth, robustness, bioengineering relevant phenotypes [9–12,14–16,23,24,97].

By integrating high-density RB-TnSeq with functional annotation, biosynthetic gene-cluster mapping, and comparative genomics, this study defines 479 essential genes in *Pseudomonas chlororaphis* ATCC 9446 and shows that essential functions are organized into coherent cellular modules, including conserved core systems for metabolism, envelope biogenesis, and information processing, rather than isolated loci. Essential modules are also embedded within biosynthetic and mobile genomic regions, revealing both deeply conserved requirements shared across *Pseudomonas* and lineage-specific constraints that are missed by current *in silico* predictors. In contrast, adaptive functions and much of secondary metabolism appear largely dispensable under the laboratory conditions tested. Notably, the recovery of multiple essential genes with unknown or poorly characterized functions underscores that some key aspects of *Pseudomonas* physiology remain unresolved. Extending this framework to additional environments and stresses will enable systematic identification of conditionally essential genes and, through barcode-based fitness profiling, move from binary essentiality toward quantitative fitness landscapes that can be integrated with physiology, targeted metabolomics, isotope tracing, and constraint-based metabolic modeling. Finally, comparative essentiality mapping across additional *Pseudomonas* species, together with the curated barcoded mutant library generated here, provides a reusable community resource to dissect gene function, regulatory networks, and metabolic integration while refining evolutionary models of minimal bacterial gene sets.

## Author statements

### Author contributions

CRediT author’s contribution: Conceptualization: A.DS., J.U. Data curation: A.DS., A.A.V.

Formal analysis: A.DS., A.A.V., J.U. Funding acquisition: J.U. Investigation: A.DS., M.A.B.G.,

J.U. Methodology: A.DS., M.A.B.G. Software: A.A.V, A.DS. Supervision: J.U. Validation: A.A.V., J.U. Visualization: A.DS., A.A.V. Writing original draft: A.DS., J.U. Writing review and editing: A.DS., A.A.V., M.A.B.G., J.U. Project administration: J.U.

## Acknowledgements

We thank Leidy Bedoya-Perez for her help in constructing the mutant library, and Gabriela Perez-Segura for technical support.

