## Supplementary Figure 1 for "Genome-wide mapping of gene essentiality in *Pseudomonas chlororaphis* ATCC 9446 using transposon mutagenesis"

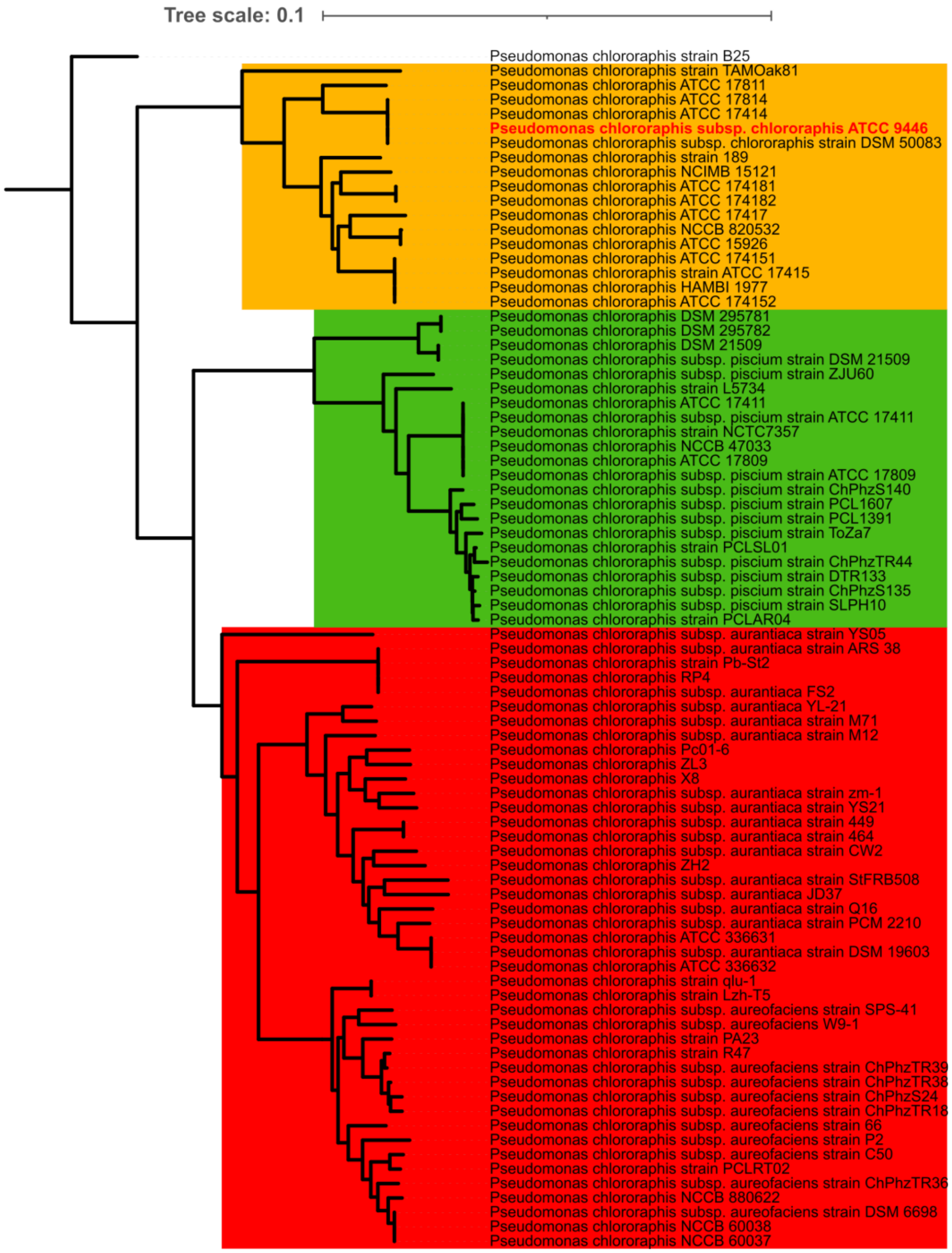


**Supplementary Figure S1.** Phylogenetic tree of the 85 *Pseudomonas chlororaphis* genomes used in this study, including Pc9446 (Red font). Major clades are highlighted by subspecies group, with yellow indicating *P. chlororaphis* subsp. *chlororaphis*, green indicating *P. chlororaphis* subsp. *piscium*, and red indicating the cluster formed by *P. chlororaphis* subsp. *aurantiaca* and *aureofaciens*. Pc9446 clustered within the subsp. *chlororaphis* group, supporting its taxonomic classification and providing phylogenetic context for the comparative analyses presented in this study.
